# Saturation kinetics and specificity of transporters for L-arginine and asymmetric dimethylarginine (ADMA) at the blood-brain and blood-CSF barriers

**DOI:** 10.1101/2025.02.14.638257

**Authors:** Mehmet Fidanboylu, Sarah Ann Thomas

## Abstract

Nitric oxide synthases (NOS) synthesize nitric oxide (NO) from L-arginine in endothelial and neuronal cells. Asymmetric dimethylarginine (ADMA) is a homologue of arginine and an endogenous inhibitor of NOS. As NO is a critical signalling molecule and influences physiological pathways in health and disease, the transfer of arginine and ADMA across the blood-CNS barriers is of interest. Our research group have previously demonstrated the presence of saturable transporters for [^3^H]-*L*-arginine and [^3^H]-ADMA at the blood-brain and blood-CSF barriers using *in vitro* and *in situ* methods. In this study, we determine the identity and kinetic characteristics of these transporters by means of the *in situ* brain/choroid plexus perfusion technique in anaesthetised mice. Results indicated that [^3^H]-arginine and [^3^H]-ADMA could be transported across blood-brain and blood-CSF barriers by the cationic amino acid transporter, system-y^+^. In contrast to the results obtained with arginine where transport was predominately by a single transport system (system-y^+^), ADMA delivery to the CNS was more complex and involved multiple transport systems (system y^+^, B^0,+^, y^+^L and b^0,+^) suggesting its concentration is tightly regulated. System y^+^ and system y^+^L transporters could be involved in the CNS to blood efflux of ADMA that we have previously observed. The half-saturation constant (K_m_) and maximal influx rate of the saturable component (V_max_) for [^3^H]-ADMA transport into the frontal cortex was 29.07±7.19 μM and 0.307±0.017 nmol.min^-1^.g^-1^, respectively, and into the CSF was 30.59±25.41 μM and 2.07±0.38 nmol.min^-1^.g^-1^, respectively. This information could help explain the arginine paradox providing evidence that ADMA interacts with transporters that can remove ADMA from cells. These removal mechanisms could be stimulated by excess arginine in the plasma resulting in increased NO production. It remains to be seen if arginine supplementation could be used to increase NO production and improve hypoperfusion observed in disease states such as Alzheimer’s and stroke.

## INTRODUCTION

There are two barriers between the blood and the central nervous system (CNS). One is called the blood-brain barrier (BBB) and is located at the brain capillary endothelial cell wall. The other is the blood-cerebrospinal fluid (CSF) barrier and is formed by the choroid plexuses and arachnoid membrane. Both barriers serve to protect the sensitive and delicate brain cells from harmful substances in the bloodstream, but at the same time allow oxygen and nutrients into the brain. The selectivity of the barriers is partly due to the presence of transporters (also called carriers) that move substances into and out of the CNS.

Over the last 30 years there have been advances in our knowledge of cationic amino acid (CAA) transport. It is now clear that there are various transporters for CAAs that show differences in site of expression, substrate specificity and mechanism. CAAs are an absolute requirement for normal brain function and yet little is known about how they enter from blood to brain in the most used laboratory model, the mouse. Of particular interest is arginine, a semi-essential CAA and an immediate precursor of nitric oxide (NO) synthesis by nitric oxide synthases (NOS). There are three isoforms of NOS including endothelial (eNOS), neuronal (nNOS) and inducible (iNOS). NO has been implicated in regulatory and mediatory roles in a variety of different physiological processes and pathological conditions involving the cardiovascular, nervous and immune systems. In particular, eNOS is critical for cardiovascular health as it plays a key role in controlling vascular tone, inhibiting inflammation and preventing thrombosis [1]. The role of NO in the brain is both complex and far reaching, and any disruption to the delicately balanced processes that control its production can be expected to have severe consequences in the brain, for example in ischaemic stroke [2]. One such molecule that is known to inhibit NO production is asymmetric dimethylarginine (ADMA) – an endogenously occurring analogue of *L*-arginine. It is believed that the inhibition of NOS by ADMA stems from the inability of NOS to utilise ADMA as a substrate [3].

In this study we will build on our previous research studies which identified saturable transporters for [^3^H]-arginine and [^3^H]-ADMA at both the blood-brain and blood-CSF barriers [4][5]. Our aim will be to explore and compare the identity of the transporters for [^3^H]-arginine and [^3^H]-ADMA at the BBB by means of the *in situ* brain perfusion technique in anaesthetised mice and the use of specific transporter inhibitors. In addition, this method allows examination of uptake directly into the CSF and so the characteristics of passage across the blood-CSF barrier, at the level of the choroid plexus, will also be investigated. We will also determine the kinetic characteristics of [^3^H]-ADMA flux into the CNS by calculation of the half-saturation constant (K_m_), maximal influx rate of the saturable component (*V_max_*) and constant of non-saturable diffusion (K_d_). Although our earlier study detected some overlap in transporter specificity by arginine and ADMA, we will test the hypothesis that as these CAAs have opposite functions, they may also use different transporters despite their structural similarities. This knowledge is essential, not only for understanding the local and neuronal physiological processes that are mediated by the CAAs these carriers transport, but also due to the widespread influence of NO in physiological pathways in health and disease and the possibility of using arginine supplementation to increase athletic performance due its indirect vasodilatory effects [6], as well as to treat endothelial cell dysfunction caused as a result of an imbalance in NO production.

## METHODS

### Materials

[^3^H]-arginine (mol. wt., 174.2 g/mol; specific activity, 43 Ci/mmol; >97% radiochemical purity) was purchased from Amersham Radiochemicals, Buckinghamshire, UK.

[^3^H]-ADMA (mol. wt., 276.7 g/mol; specific activity, 8 Ci/mmol; 96.4% radiochemical purity) was synthesised and tritiated by Amersham Radiochemicals, Cardiff, UK.

[^14^C]-sucrose (mol. wt., 342.3 g/mol; specific activity, 0.412 Ci/mmol; 99% radiochemical purity; Moravek Biochemicals, Brea, CA, USA). Unlabelled *L*-arginine and asymmetric dimethylarginine (N^G^,N^G^-dimethylarginine dihydrochloride or ADMA) were purchased from Sigma Aldrich (Dorset, UK). All transport inhibitors used were purchased from Sigma Aldrich; Dorset, UK and were readily soluble in the artificial plasma – requiring no extra steps for dissolution beyond standard mechanical stirring and heating to physiological temperature (37 °C).

### Animals

All experiments requiring the use of animals were performed in accordance with the Animal (Scientific Procedures) Act, 1986 and Amendment Regulations 2012 and with consideration to the Animal Research: Reporting of *In Vivo* Experiments (ARRIVE) guidelines. The study was approved by the King’s College London Animal Welfare and Ethical Review Body and performed under license: 70/6634. In the following sections we also describe attempts made to reduce the number of animals used in our studies in line with the 3Rs (replacement, refinement, and reduction). It is noted that all the data was obtained during the same time-frame by the same researcher.

All animals used in procedures were adult male BALB/c mice (between 23 g and 25 g) sourced from Harlan Laboratories, Oxon, UK, unless otherwise stated. All animals were maintained under standard temperature and lighting conditions. Access to water and food was provided *ad libitum*. Welfare was monitored daily by animal technicians. Animals which were showing signs of distress (such as not feeding or drinking) were brought to the attention of the named veterinary surgeon and the animal euthanized. Mice were used for all the procedures under a non-recovery anaesthetic and represent the animals of lowest neurological sensitivity to which the protocols can be successfully applied. Domitor® (medetomidine hydrochloride) and Vetalar® (ketamine) were both purchased from Harlan Laboratories; Cambridge, UK. All animals were anaesthetised (2 mg/kg Domitor® and 150 mg/kg Vetalar® injected intraperitoneally), and a lack of self-righting and paw-withdrawal reflexes (as surrogate indicators of consciousness) thoroughly checked prior to carrying out all procedures. Animals were heparinised with 100 units heparin (in 0.9% m/v NaCl(aq), Harlan Laboratories; Oxon, UK), administered *via* the intraperitoneal route prior to surgery.

### *In situ* brain perfusion technique

The *in situ* brain /choroid plexus perfusion method was performed as previously described and will only be briefly detailed here [7][5]. An artificial plasma containing [^3^H]-arginine (11.6 nM) or [^3^H]-ADMA (62.5 nM) and [^14^C]-sucrose was infused into the left ventricle of the heart for 10 minutes at a flow rate of 5 mL/min. The artificial plasma consisted of a modified Krebs-Henseleit mammalian Ringer solution with the following constituents: 117 mM NaCl, 4.7 mM KCl, 2.5 mM CaCl_2_, 1.2 mM MgSO_4_, 24.8 mM NaHCO_3_, 1.2 mM KH_2_PO_4_, 10 mM glucose, and 1 g/liter bovine serum albumin. At the end of the perfusion period a CSF sample was taken from the cisterna magna, the animal was decapitated, the brain removed and samples taken under a Leica S4E L2 stereomicroscope (Leica; Buckinghamshire, UK). The samples included the frontal cortex, caudate nucleus, occipital cortex, hippocampus, hypothalamus, thalamus, pons and cerebellum. The circumventricular organs (CVOs) including the choroid plexus, pituitary gland and pineal gland were also sampled. Brain samples were also taken for capillary depletion analysis.

[^14^C]-sucrose is a baseline marker molecule. It is a polar, hydrophilic molecule, which does not cross membranes well, and can be used to measure extracellular space including the vascular space. It is specifically used in brain samples (including brain homogenate and brain supernatant) to measure the vascular space at all perfusion time points. At later time points, in particular, it may also represent the capillary endothelial cell volume and / or diffusion across the brain capillary wall and into the brain interstitial fluid. In the pellet samples it will measure extracellular fluid and may also represent capillary endothelial cell volume. In the choroid plexus samples, [^14^C]-sucrose would represent vascular space, plus the interstitial fluid volume in the compartment between choroid plexus capillary endothelial cells and the choroid plexus epithelial cells of the blood-CSF barrier. It may also represent any accumulation into the cells of the choroid plexus. In the pituitary and pineal gland samples, [^14^C]-sucrose would measure the vascular space plus the interstitial fluid compartment between the blood vessels and the tanycytic barrier. In CSF samples, [^14^C]-sucrose can be used to correct for possible contamination with blood.

#### Capillary depletion analysis

Capillary depletion analysis involved preparing a whole brain homogenate by the addition of a capillary depletion buffer (141 mM NaCl, 4.0 mM KCl, 2.8 mM CaCl_2_, 1.0 mM MgSO_4_, 10.9 mM HEPES, 1.0 mM NaH_2_PO_4_, 10 mM glucose at pH 7.4) and a dextran (MW 60,000-90,000) solution (final concentration 13%) [7][5]. Both the physiological capillary depletion buffer and dextran solution were maintained at 4 °C both to halt any cell metabolism and transport processes, and to reduce any cellular or protein damage due to heat generated from the homogenisation process. The whole brain homogenate was then taken for dextran density centrifugation (5,400 x g) and 4 °C to produce an endothelial cell-enriched pellet and supernatant containing brain parenchyma.

#### Liquid scintillation analysis

All samples (brain regions, brain homogenate, supernatant, pellet, CVOs, CSF and plasma samples) were solubilised in 0.5 mL tissue solubiliser (Solvable; Perkin-Elmer; Boston, MA, USA). After incubating the samples at room temperature for 48 h, 4 mL scintillation fluid (Lumasafe®; Perkin-Elmer; Boston, MA, USA) was added to each before vigorous vortexing. The amount of [^3^H]- and [^14^C]-radioactivity in each sample were then quantified using a Packard Tri-Carb 2900TR liquid scintillation counter (Perkin-Elmer; Boston, MA, USA). Counts per minute were then converted to disintegrations per minute (dpm) by the counter using internally stored quench curves from standards.

### Experimental design

#### Transporter specificity experiments

The identity of the transporter(s) was studied by measuring the [^3^H]-arginine or [^3^H]-ADMA uptake into the CNS in the absence and presence of specific transporter inhibitors in the artificial plasma at a perfusion time of 10 minutes and the results compared. These inhibitors included 20 mM *L*-homoarginine, 4 mM 2-amino-endo-bicyclo[2.2.1]heptane-2-carboxylic acid (BCH), 500 μM a-methyl-*D*,*L*-tryptophan, 200 μM *L-*phenylalanine, 5 mM *L*-leucine and 2 mM harmaline. The transporter proteins these inhibitors target are listed in S1 Table.

It is noted that the data obtained from the [^3^H]-arginine control group at 10 minutes and the [^3^H]-ADMA control group at 10 minutes data have recently been published [5]. In this present study, these data sets were combined with new data sets obtained using the transporter inhibitors to identify specific transporter interaction. This allowed us to reduce the number of animals needed for this study in line with the 3Rs (replacement, refinement, and reduction).

#### Self-inhibition experiments

A series of single-time point experiments involved varying the unlabelled concentration of ADMA in the artificial plasma and measuring the [^3^H]-ADMA uptake into the brain regions, capillary depletion samples, CVOs and CSF after 10 min of perfusion. Specifically, the uptake of [^3^H]-ADMA (62.5 nM) and [^14^C]-sucrose into the CNS was measured in the absence and presence of 0.5, 3.0, 10, 100 and 500 μM un-labelled ADMA in the artificial plasma for a perfusion period of 10 minutes. This enabled calculation of the kinetic constants; *V*_max_ (the maximal influx rate of the saturable component), *K*_m_ (the half-saturation constant) and *K*_d_ (the constant of non-saturable diffusion).

These concentrations were selected as the plasma concentration for ADMA under normal physiological conditions is typically found to be approximately 0.5 μM in humans and in several mammalian species, including the mouse [8–11]. Plasma concentrations for ADMA under pathological conditions are typically in the range of 3.0 μM [12,13]. Concentrations above 10 μM can be considered as supraphysiological as ADMA would not ordinarily be found at concentrations that high in the body in health or disease. However, the higher ADMA concentrations can help determine the kinetic characteristics of transport.

It is noted that the data obtained from the [^3^H]-ADMA group in the presence of 100 μM unlabelled ADMA data has recently been published[5]. In this present study this existing data set was combined with new data sets using unlabelled ADMA at four other concentrations (0.5, 3.0, 10 and 500 μM) in order to calculate previously unreported kinetic constants (K_m_, V_max_ and K_d_). This allowed us to reduce the number of animals needed in line with the 3Rs. Kinetic constants are valuable as they allow data from different assays to be compared.

### Expression of results

#### Relative uptake from the perfusate into the tissues

The concentration of radioactivity in the brain regions (C_Brain_ ; dpm/g), CVOs (C_CVO_ ; dpm/g) and CSF (C_CSF_ ; dpm/ml) is expressed as a percentage of that in the artificial plasma (C_pl_; dpm/ml) and termed *R_BRAIN_*(dpm/100g), *R_CVO_* (dpm/100g) or *R_CSF_* (dpm/100ml) respectively or R_Tissue_ as shown in equation 1.

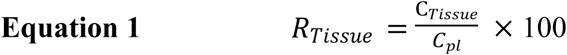

Correcting for vascular space involved subtracting the *R_Tissue_* value obtained for [^14^C]-sucrose in each sample from the *R_Tissue_* value concurrently obtained for the [^3^H]-labelled solute of interest (i.e. [^3^H]-arginine or [^3^H]-ADMA).

The percentage change in the R_Tissue_ uptake values achieved in the absence or presence of an unlabelled inhibitor can be determined by means of equation 2.

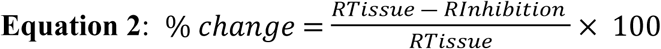

Where R_Inhibition_ is the R_Tissue_ uptake in the presence of an inhibitor in the artificial plasma. A decrease in the uptake of radiolabelled solute in the presence of unlabelled inhibitor is indicative of a saturable influx transport system. Conversely, an increase in the uptake of radiolabelled solute in the presence of unlabelled inhibitor is indicative of a saturable efflux transport system. No change in the distribution of the radiolabelled solute in the absence and presence of the unlabelled inhibitor may indicate the absence of saturable transport by the radiolabelled solute or the use of influx and efflux transporters by the radiolabelled solute.

#### Calculation of kinetic constants

The rate of transfer across the BBB and the blood-CSF barrier can be calculated as a transfer constant (K_in_) as shown in equation 3 [5].

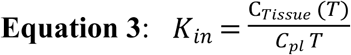

Where C_Tissue_ (T) is radioactivity (dpm) per g of tissue at time-point T (perfusion time in minutes), and C_pl_ is radioactivity (dpm) per mL of artificial plasma.

This equation assumes that the entry of the radiolabelled solute of interest into the CNS is proportional to, but less than, its concentration in the artificial plasma and efflux (CNS to blood) is much smaller than influx (blood to CNS) of the test solute and therefore can be ignored [14].

It should however be noted that calculating the K_in_ value from blood to CNS using this method requires that *R_Tissue_* at time T is first corrected for vascular space by subtracting the *R_Tissue_* value for [^14^C]-sucrose determined at that time-point.

In the case where the uptake of the radiolabelled test amino acid is examined in the presence of various unlabelled concentrations of the same test solute, K_in_, can be defined as in equation 4:

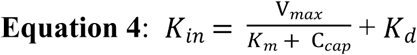

Where V_max_ is the maximal influx rate of the saturable component; K_m_ is the half-saturation constant; K_d_ is the constant of non-saturable diffusion and C_cap_ is the mean capillary concentration of the amino acid.

Under the present experimental conditions, flow to the brain (F) is always greater than 1 ml.min^-1^.g^-1^, which is much greater than the highest measured K_in_ (i.e. F >> K_in_), the difference between C_Cap_ and concentrations of amino acid in the artificial plasma, C_pl_, becomes negligible and equation 4 can be simplified to equation 5 :

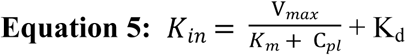

Unidirectional ADMA flux (J_in_) into the brain (nmol.min^-1^.g^-1^) and CSF (nmol.min^-1^.ml^-1^) was determined using equations 6 and 7:

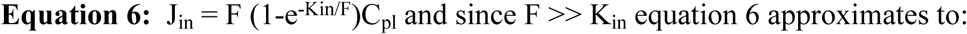

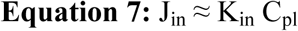

Unidirectional influx of amino acids (J_in_) into the CNS can be related to the kinetic constants K_m_, V_max_ and K_d_ by equation 8 where total flux is equal to the sum of the saturable flux and non-saturable flux:

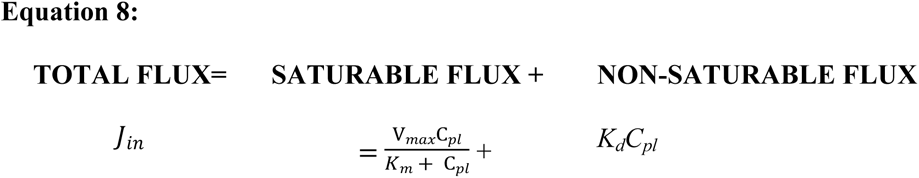

When appropriate, K_d_ was determined as the slope of the line from linear regression analysis of the total flux and ADMA concentration at the highest two concentrations utilized (100 and 500 μM). Linear regression analysis of the data was performed using Graph Pad Prism (Version 10.2.2). This constant of non-saturable diffusion was then used to calculate a non-saturable flux which could be subtracted from the total flux to determine the saturable flux. Saturable flux was then plotted as a function of concentration and using Michaelis-Menten Kinetic analysis in Graph Pad Prism (Version 10.2.2) estimates of the best-fit values of V_max_ and K_m_ obtained. In some cases, there was no measurable non-saturable component and so estimates of V_max_ and K_m_ were obtained from plots of total flux, which was equal to saturable flux.

### Statistics

Data from all experiments are presented as mean ± standard error of the mean (SEM). The samples were grouped into brain regions, capillary depletion samples and CVOs and CSF for presentation and data analysis as appropriate. Statistical significance was taken as follows: not significant (ns; *p* > 0.05) or significant (\**p* < 0.05, \*\**p* < 0.01, \*\*\**p* < 0.001). Statistical analyses were performed using GraphPad Prism v5.0c or v6 graphing and statistics package for Mac or using GraphPad Prism v10.2.2 for Windows.

## RESULTS

### Arginine

The transport of [^3^H]-arginine and [^14^C]-sucrose across the blood-brain and blood-CSF barriers was examined in the presence of transporter inhibitors. All inhibitors were used at previously published concentrations known to inhibit specific transport systems (S1 Table). [^14^C]-sucrose distribution into each of the brain regions, capillary depletion samples, CVO and CSF was not significantly affected by the presence of 20 mM *L*-homoarginine, 4 mM BCH or 500 μM α-methyl-*D*,*L-*tryptophan (S1-S3 Figs). This indicates that these inhibitors did not affect the integrity of the blood-brain and blood-CSF barriers in the [^3^H]-arginine experiments.

The uptake of [^3^H]-arginine is almost totally inhibited (up to 99.7%) by 20 mM *L*-homoarginine in all brain regions, except the hypothalamus where the reduced uptake of 98.0% did not attain statistical significance due to the wider variability of the values in this region (Figure 1). The inclusion of 4 mM BCH or 500 μM α-methyl-*D*, *L*-tryptophan in the artificial plasma did not significantly affect the uptake of [^3^H]-arginine uptake in the majority of brain regions sampled. The exception was the hippocampus sample where the inclusion of 500 μM α-methyl-*D*, *L*-tryptophan in the artificial plasma significantly inhibited [^3^H]-arginine uptake by 49.4%.

**Figure 1:**
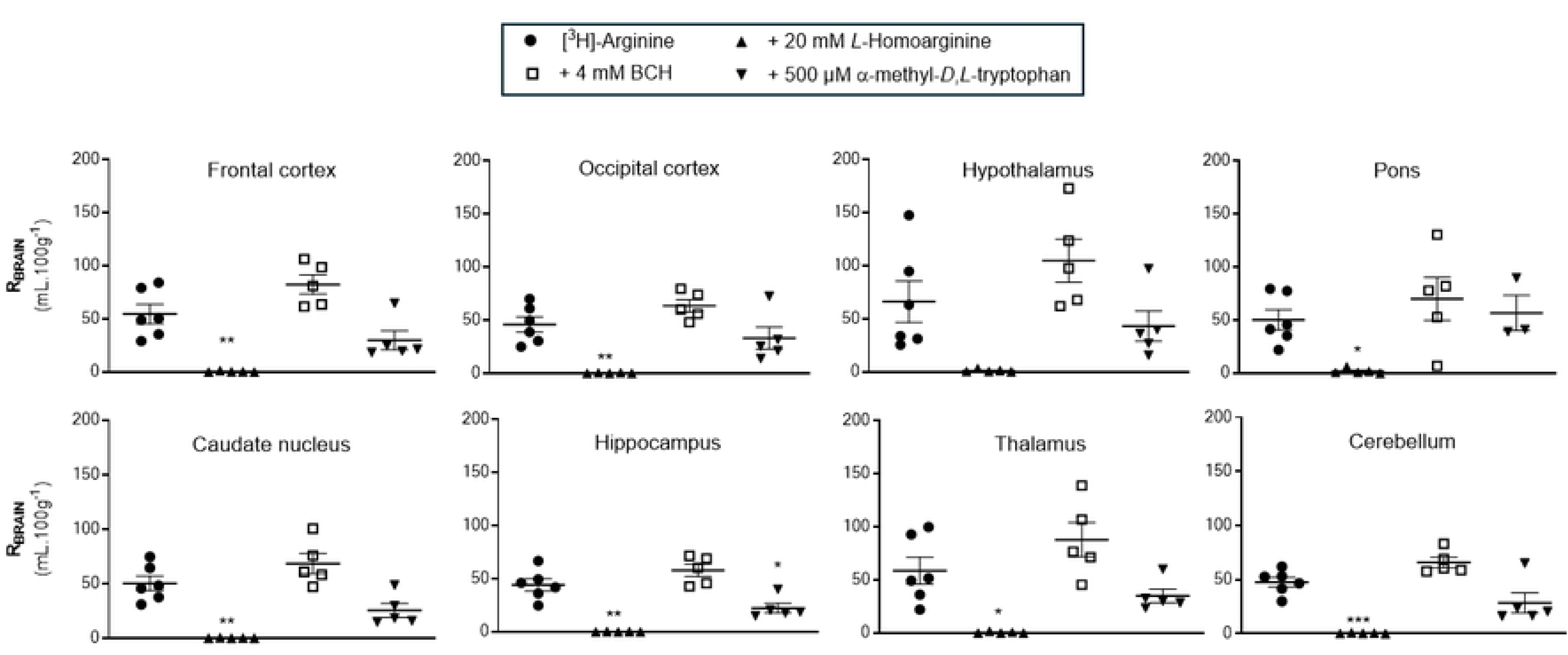
The effect of *L*-homoarginine, BCH and α-methyl-*D*,*L*-tryptophan on the regional brain uptake of [^3^H]-arginine (10 minute perfusion). Uptake is expressed as the percentage ratio of tissue to plasma (mL.100 g^-1^) and is corrected for [^14^C]-sucrose (vascular space). Each bar represents the mean ± SEM of 5-6 animals, except pons where it was n of 3 for the α-methyl-*D,L*-tryptophan group. Each marker represents one animal. One-way ANOVA with Dunnett’s post-hoc test was used to compare means to control ([^3^H]-arginine alone), with statistical significance taken as \**p* <0.05, **p<0.01, ***p<0.001 (GraphPad Prism 10.2 for Windows).

As you may expect from the data described above, the uptake of [^3^H]-arginine was also almost totally inhibited (up to 99.7%) by 20 mM *L*-homoarginine in the whole brain homogenate, supernatant containing brain parenchyma, and endothelial cell-enriched pellets following capillary depletion analysis (Figure 2). There was also no significant effect of 4 mM BCH or 500 μM α-methyl-*D*,*L-*tryptophan on [^3^H]-arginine distribution in these samples.

**Figure 2:**
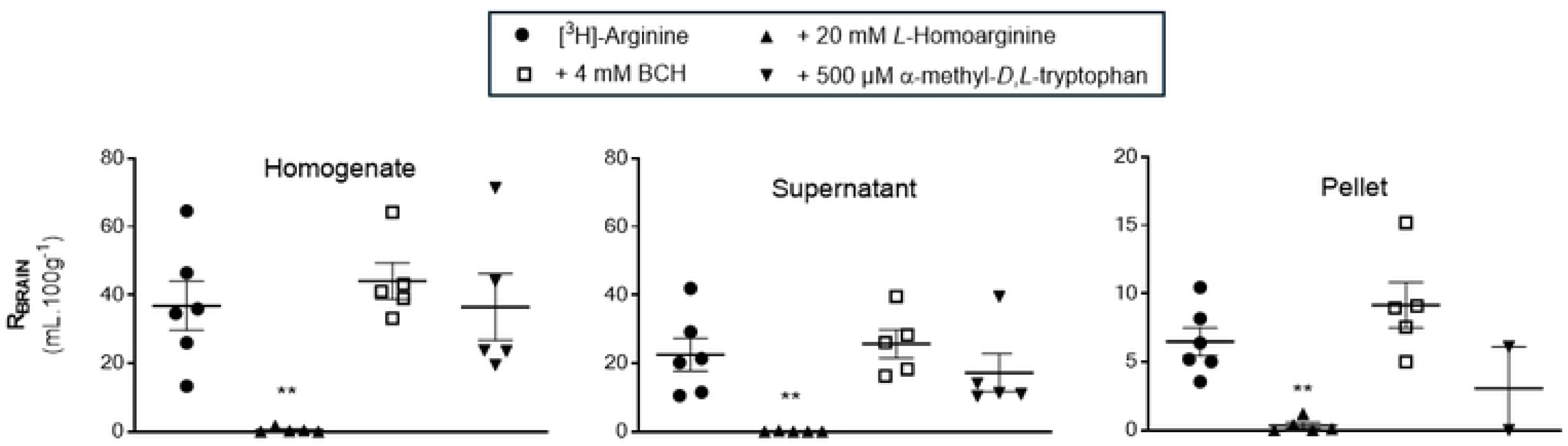
The effect of *L*-homoarginine, BCH and α-methyl-*D*,*L*-tryptophan on the distribution of [^3^H]-arginine in capillary depletion samples (10 minute perfusion). Uptake is expressed as the percentage ratio of tissue to plasma (mL.100 g^-1^) and is corrected for [^14^C]-sucrose. Each bar represents the mean ± SEM of 4-6 animals, except the α-methyl-*D*,*L*-tryptophan pellet group where it was n=2. Each marker represents one animal. One-way ANOVA with Dunnett’s post-hoc test was used to compare means to control ([^3^H]-arginine alone), with statistical significance taken as \**p* <0.05, **p<0.01, ***p<0.001 (GraphPad Prism 10.2.2 for Windows).

The distribution of [^3^H]-arginine into the choroid plexus and pituitary gland was significantly inhibited by 20 mM *L*-homoarginine being 99.8% and 97.6% respectively (Figure 3). Interestingly, although there was a decrease of 88.4% in the distribution of [^3^H]-arginine in the presence of *L*-homoarginine into the CSF, this failed to attain statistical significance. This will be related to the difficulties in taking CSF samples in these small animals and as a consequence the low sample size of 3 for this inhibitor group (Figure 3). In agreement with observations in all the other samples, (except the hippocampus), the inclusion of either 4 mM BCH or 500 μM α-methyl-*D*,*L-*tryptophan in artificial plasma had no statistically significant effect on the distribution of [^3^H]-arginine into the CSF, choroid plexus and pituitary gland.

**Figure 3:**
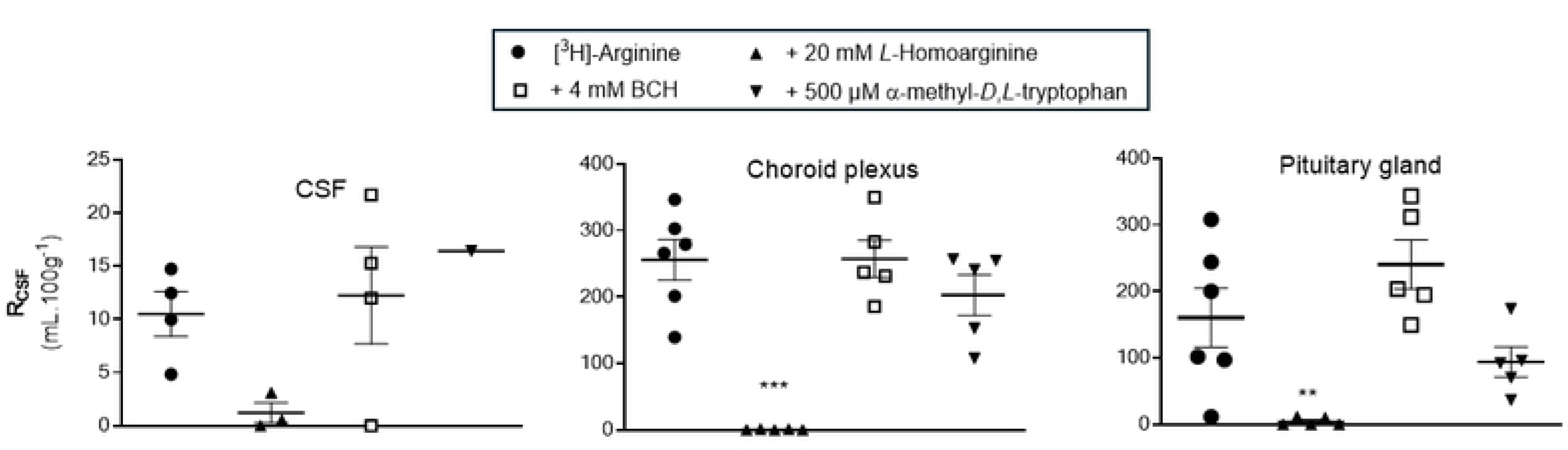
The effect of *L*-homoarginine, BCH and α-methyl-*D*,*L*-tryptophan on the distribution of [^3^H]-arginine in CSF, choroid plexus and pituitary gland (10 minute perfusion). Uptake is expressed as the percentage ratio of tissue or CSF to plasma (mL.100 g^-1^) and is [^14^C]-sucrose corrected. Each bar represents the mean ± SEM of 3-6 animals except the α-methyl-*D*,*L*-tryptophan CSF group where it was n=1. Each marker represents one animal. One-way ANOVA with Dunnett’s post-hoc test was used to compare means to control ([^3^H]-arginine alone), with statistical significance taken as \**p* <0.05, **p<0.01, ***p<0.001 (GraphPad Prism 10.2.2 for Windows).

#### ADMA

The transport of [^3^H]-ADMA and [^14^C]-sucrose across the blood-brain and blood-CSF barriers was also examined in the presence of specific transport inhibitors (S1 Table). All inhibitors were used at previously published concentrations known to inhibit the specific transport systems of interest. To explore if the inhibitor affected the integrity of the blood-CNS interfaces we first compared the [^14^C]-sucrose values in the absence and presence of the various inhibitors in these [^3^H]-ADMA experiments (S4-S6 Fig).

[^14^C]-sucrose distribution into each of the brain regions including the capillary depletion analysis compartments and CVOs was not significantly affected by most of the inhibitors i.e. 20 mM *L*-homoarginine, 4 mM BCH, 500 μM α-methyl-*D*,*L-*tryptophan or 2 mM harmaline (S4-S6 Fig). However, two of the inhibitors, 5 mM *L*-leucine and 200 μM *L*-phenylalanine did significantly change [^14^C]-sucrose distribution in some of the samples. Specifically, the [^14^C]-sucrose space was significantly higher than expected with 5 mM *L*-leucine in the following samples: frontal cortex, caudate nucleus, cerebellum and pellet (S4 Fig and S5 Fig). This suggests that L-leucine at a concentration of 5 mM could damage membranes (pellet; S5 Fig) resulting in loss of BBB integrity (brain regions, S4 Fig). Interestingly, 200 μM *L*-phenylalanine caused a significant reduction in the vascular space of the occipital cortex and hippocampus possibly as a result of vasoconstriction and/or inadequate perfusion (S4 Fig). However, it is noted that the [^14^C]-sucrose values achieved with phenylalanine in these two regions are well within the range of [^14^C]-sucrose values achieved with the other inhibitor groups (S4 Fig), so the difference observed with phenylalanine could be ignored. Although in these experiments a reduction of vascular space is less of a concern than an increase in vascular space, which suggests loss of BBB integrity, the effects of both these inhibitors on the distribution of [^3^H]-ADMA into these specific samples were not used to draw any conclusions about the transporter sensitivity of [^3^H]-ADMA. However, all data sets are presented for comparison (Figures 4 and 5; S2 and S3 Tables).

**Figure 4:**
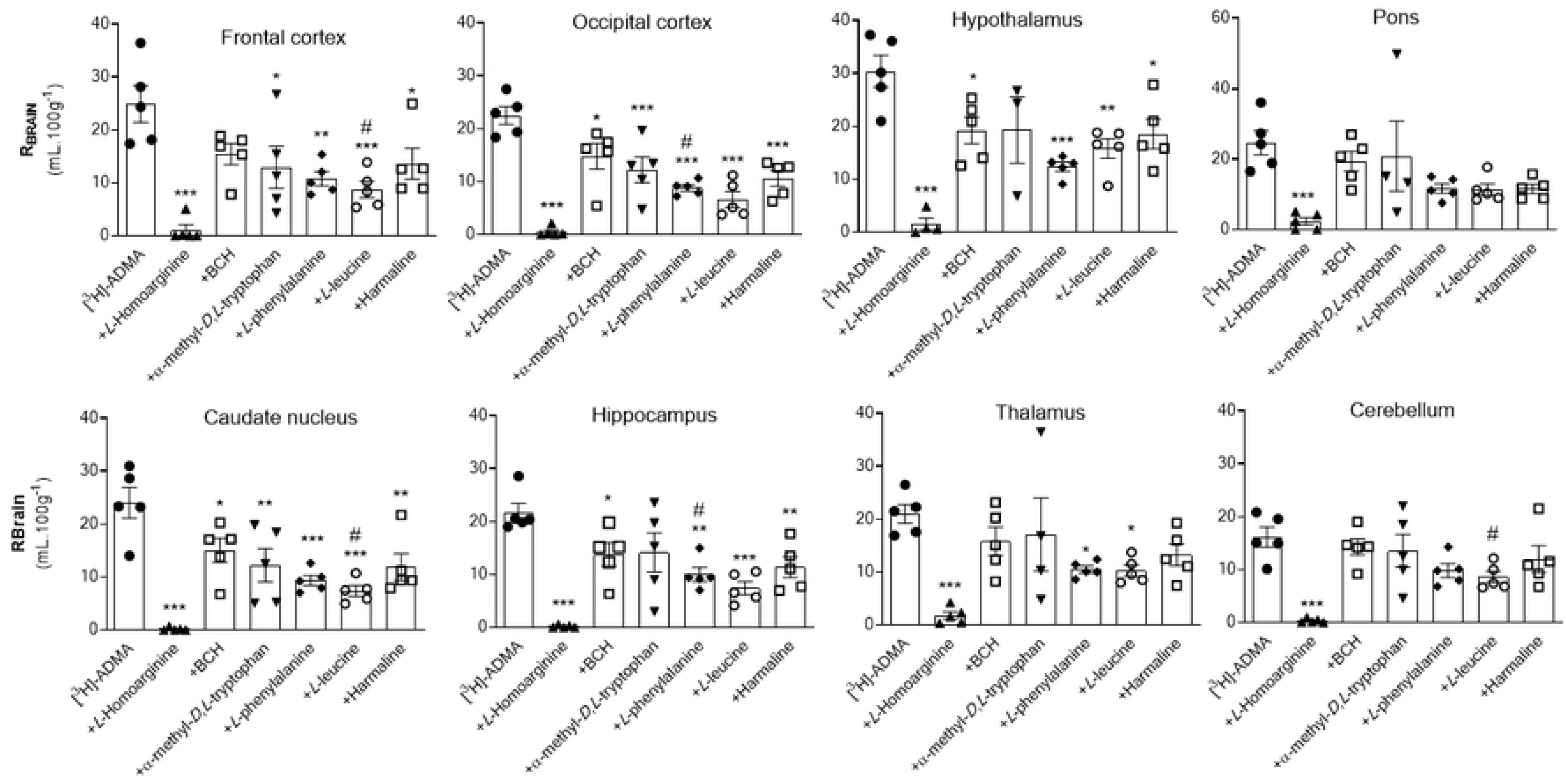
The effect of 20 mM *L*-homoarginine, 4 mM BCH, 500 μM α-methyl-*D*,*L*-tryptophan, 200 μM *L-* phenylalanine, 5 mM *L*-leucine and 2 mM harmaline on the regional brain uptake of [^3^H]-ADMA (10 minute perfusion). Uptake is expressed as the percentage ratio of tissue to plasma (mL.100 g^-1^) and is corrected for [^14^C]-sucrose (vascular space). Perfusion time is 10 minutes. Each bar represents the mean ± SEM of 4-5 animals. Each marker represents one animal. Asterisks represent one-way ANOVA with Dunnett’s post-hoc tests comparing mean±SEM to control within each sample/region, \**p* < 0.05, \*\**p* < 0.01, \*\*\**p* < 0.001 (GraphPad Prism 6.0 for Mac). **^#^**The [^14^C]-sucrose (vascular space; S4 Figure) in that brain region was statistically affected by the presence of the inhibitor (i.e. *L-*phenylalanine or *L-*leucine).

**Figure 5:**
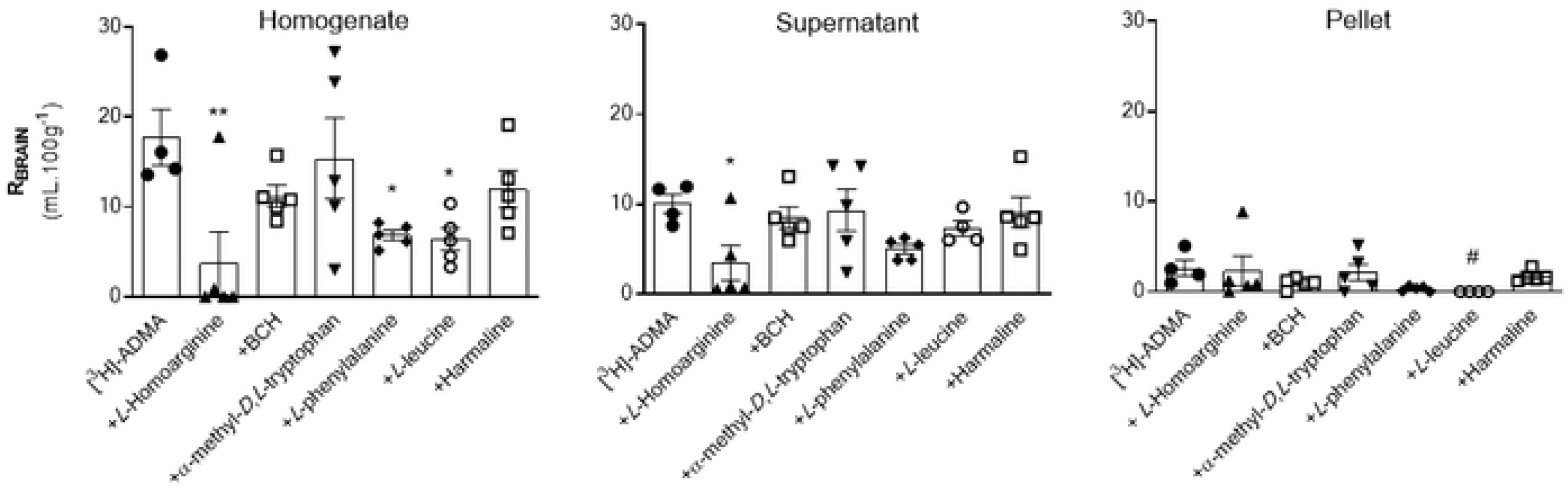
The effect of 20 mM *L*-homoarginine, 4 mM BCH, 500 μM α-methyl-*D*,*L*-tryptophan, 200 μM *L-* phenylalanine, 5 mM *L*-leucine and 2 mM harmaline on the distribution of [^3^H]-ADMA in capillary depletion samples (10 minute perfusion). Uptake is expressed as the percentage ratio of tissue to plasma (mL.100 g^-1^) and is corrected for [^14^C]-sucrose. Perfusion time is 10 minutes. Each bar represents the mean ± SEM of 4-5 animals. Each marker represents one animal. Asterisks represent one-way ANOVA with Dunnett’s post-hoc tests comparing mean±SEM to control within each region/sample, \**p* < 0.05, \*\**p* < 0.01 (GraphPad Prism 6.0 for Mac). **^#^**The [^14^C]-sucrose value (S5 Figure) in the pellet was statistically affected by the presence of L-leucine.

The uptake of [^3^H]-ADMA is almost totally inhibited by 20 mM *L*-homoarginine in all brain regions (up to 99.2%; Figure 4; S2 Table). The inclusion of either 4 mM BCH, 500 μM α-methyl-*D*,*L*-tryptophan, 200 μM *L*-phenylalanine, 5 mM *L*-leucine, or 2 mM harmaline in artificial plasma also reduced [^3^H]-ADMA uptake into brain regions, but to a much lesser degree, and only attaining statistical significance in some regions (Figure 4; S2 Table). It is noted that the inhibitors, L-leucine and L-phenylalanine, could still significantly inhibit [^3^H]-ADMA uptake in those brain regions which still had an intact BBB as measured by [^14^C]-sucrose (e.g. hypothalamus and thalamus; S2 Table). *L*-homoarginine also inhibited uptake of [^3^H]-ADMA in whole brain homogenate (by 78.9%) and brain parenchyma (supernatant; by 65.8%) following capillary depletion (Figure 5; S3 Table). *L*-phenylalanine and *L*-leucine inhibited [^3^H]-ADMA distribution into whole brain homogenate (by 61.3% and 63.7% respectively), but not brain parenchyma (supernatant) following capillary depletion (Figure 5; S3 Table). The inclusion of BCH, α-methyl-*D*,*L*-tryptophan or harmaline into the artificial plasma did not significantly affect the uptake of [^3^H]-ADMA into either the brain homogenate or supernatant. None of the inhibitors inhibited uptake of [^3^H]-ADMA in the endothelial cell enriched pellet (Figure 5; S3 Table).

None of the inhibitors affected the distribution of [^3^H]-ADMA in CSF (Figure 6; S4 Table). All the inhibitors inhibited the distribution of [^3^H]-ADMA in the pineal gland (Figure 6; S4 Table). The distribution of [^3^H]-ADMA in the choroid plexus and pituitary gland was inhibited by all inhibitors except for 500 μM α-methyl-*D,L*-tryptophan (Figure 6; S4 Table).

**Figure 6:**
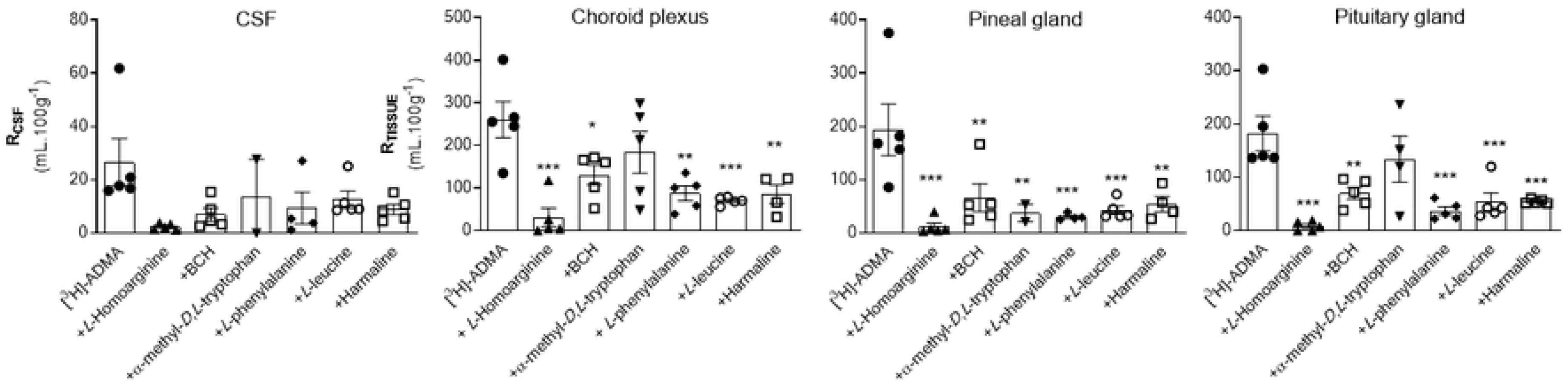
The effect of 20 mM *L*-homoarginine, 4 mM BCH, 500 μM α-methyl-*D*,*L*-tryptophan, 200 μM *L-* phenylalanine, 5 mM *L*-leucine and 2 mM harmaline on the distribution of [^3^H]-ADMA in CSF, choroid plexus and CVOs (10 minute perfusion). Uptake is expressed as the percentage ratio of tissue or CSF to plasma (mL.100 g^-1^) and has been corrected for [^14^C]sucrose. Perfusion time is 10 minutes. Each bar represents the mean ± SEM of 4-5 animals except for the CSF and pineal sample where with the inhibitor group, α-methyl-*D,L*-tryptophan n = 2. Each marker represents one animal. Asterisks represent one-way ANOVA with Dunnett’s post-hoc tests comparing mean±SEM to control within each region/sample, \**p* < 0.05, \*\**p* < 0.01, \*\*\**p* < 0.001 (GraphPad Prism 6.0 for Mac).

### Kinetic constants

The transport of [^3^H]-ADMA (62.5 nM) and [^14^C]-sucrose across the blood-brain and blood-CSF barriers was examined in the presence of different concentrations of unlabelled ADMA (0.5-500 μM) and this data was used to calculate the kinetic constants, K_m_, V_max_ and K_d_. To explore if the unlabelled ADMA affected the integrity of the blood-CNS interfaces we first compared the [^14^C]-sucrose values in the absence and presence of the various concentrations of unlabelled ADMA (S7-S9 Fig).

The [^14^C]-sucrose distribution into the brain regions, brain homogenate, brain supernatant, CSF and CVOs in the presence of unlabelled ADMA up to a concentration of 100 μM were not statistically significantly different to the control values achieved in the absence of unlabelled ADMA (S7-S9 Fig). The exception to this is the [^14^C]-sucrose values in the presence of 3 μM unlabelled ADMA, which were statistically different to the control values achieved in the absence of unlabelled ADMA in two brain regions, the hippocampus and thalamus (S7 Fig). However, we considered that this difference with 3 μM unlabelled ADMA could be ignored as the values were within the same range as that achieved for [^14^C]-sucrose in the presence of the higher unlabelled ADMA concentrations where no statistical difference was observed (i.e. 10 and 100 μM unlabelled ADMA). In contrast, unlabelled ADMA concentrations of 500 μM did statistically increase the distribution of [^14^C]-sucrose into the occipital cortex, caudate nucleus, hippocampus, thalamus, pons and cerebellum and the values measured were at the upper range of [^14^C]-sucrose values achieved at the lower concentrations of unlabelled ADMA (i.e. ≤100 μM unlabelled ADMA; S7 Fig). As this does suggest loss of BBB integrity in these regions we only interpreted the kinetic characteristics of [^3^H]-ADMA in the other regions where the BBB remained statistically intact (i.e. frontal cortex, hypothalamus, homogenate and supernatant) (S7 Fig and S8 Fig). Interestingly, the [^14^C]-sucrose distribution into the pellet samples was statistically reduced by the presence of unlabelled ADMA at most concentrations (S8 Fig) and was in the range achieved in other test groups where no statistical difference was obtained (S2 Fig). This suggests the membrane was intact in these samples and we could present and interpret the kinetic characteristics of ADMA distribution in this sample as well.

The uptake of [^3^H]-ADMA (62.5 nM) was significantly self-inhibited by 0.5, 3.0, 10, 100 and 500μM un-labelled ADMA in all brain regions (S10 Fig). The uptake of [^3^H]-ADMA into the capillary depletion samples, pineal gland, pituitary gland and choroid plexuses was decreased in the presence of unlabelled ADMA, although it failed to attain statistical significance in the homogenate, supernatant and pellet at a concentration of 10 μM unlabelled ADMA and supernatant at a concentration of 0.5 μM unlabelled ADMA (S11-S12 Figs). Distribution of [^3^H]-ADMA into the CSF was significantly decreased at all concentrations of unlabelled ADMA except the lowest concentration of 0.5 μM, where it failed to reach statistical significance (S12 Fig).

The R_Tissue_ uptake data for [^3^H]-ADMA in the absence and presence of unlabelled ADMA into the brain regions, capillary depletion samples and CSF shown in S10-S12 Fig, has been [^14^C]-sucrose corrected, and was used to calculate the total flux of [^3^H]-ADMA. The total flux, saturable flux and non-saturable flux of [^3^H]-ADMA into the CNS have been plotted as a function of ADMA plasma concentration and are shown in Figure 7, Figure 8 and S13 Fig. The non-saturable fluxes (K_d_C_pl_) have been determined from linear regression analysis as described in the methods. The saturable flux data had been calculated by subtracting the non-saturable flux from the total flux, where appropriate. The total flux of [^3^H]-ADMA into the CSF was equal to the saturable flux and therefore there was no non-saturable component (Figure 8). The K_m_ and the V_max_ were derived using non-linear regression analysis of the saturable flux data (Enzyme kinetics – Michaelis-Menten, GraphPad Prism 10.0 for Windows). Table S5 shows the values for K_m_, V_max_ and K_d_. Interestingly, the K_m_, V_max_ and K_d_ values in each brain region are similar, even if there was loss of BBB integrity in that region as measured by [^14^C]-sucrose (Figures S13; S5 Table).

**Figure 7:**
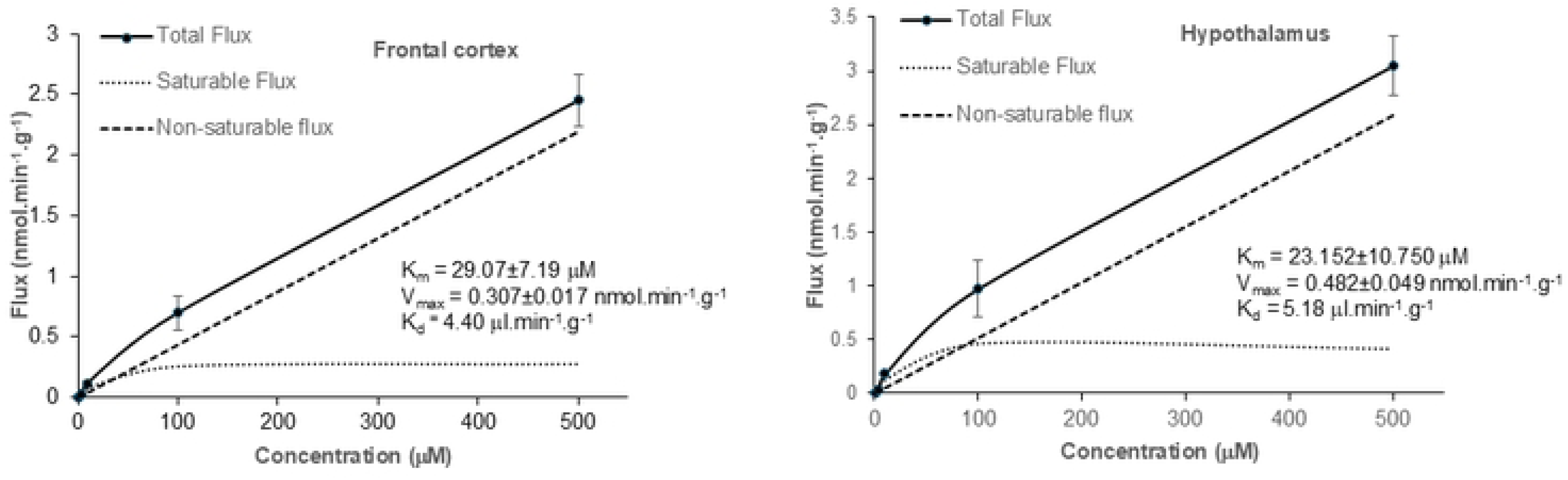
The contributions of the saturable and non-saturable components to total brain flux of [^3^H]-ADMA are plotted against the unlabelled ADMA concentration. The measured values are the mean ±SEM for 4-5 mice at each of the 6 ADMA concentrations and has been [^14^C]-sucrose corrected. The lower lines show the contributions of the saturable and non-saturable components to total influx. The K_d_ value was calculated from linear regression analysis of the total flux at the highest concentrations at 100 and 500 μM. The K_m_ and V_max_ were calculated by Michaelis-Menten kinetic analysis of the saturable flux (mean values were used). Analyses were performed using GraphPad Prism version10.

**Figure 8:**
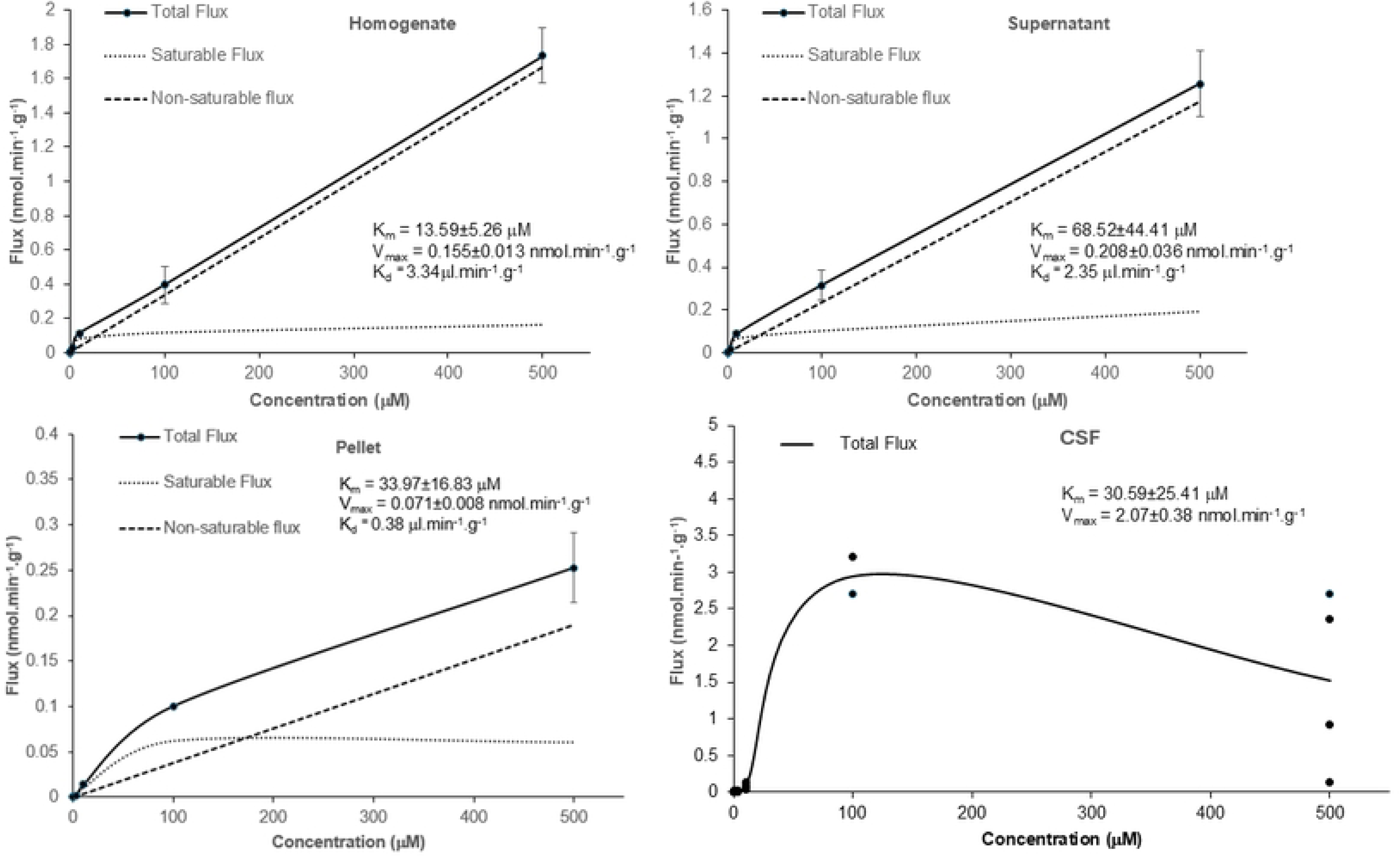
The contributions of the saturable and non-saturable components to total flux of [^3^H]-ADMA into the capillary depletion samples are plotted against the unlabelled ADMA concentration. The measured values are the mean ±SEM for 4-5 mice (homogenate), 4-5 mice (supernatant) and 5 mice (pellet except at 100 μM where it was 1 mouse) at each of the 6 ADMA concentrations and has been [^14^C]-sucrose corrected. The lower lines show the contributions of the saturable and non-saturable components to total influx. The K_d_ value was calculated from linear regression analysis of the total flux at 100 and 500 μM. The K_m_ and V_max_ were calculated by Michaelis-Menten kinetic analysis of the saturable flux (mean values were used). Analyses were performed using GraphPad Prism version10. **The total flux of [^3^H]-ADMA into the CSF is plotted against the unlabelled ADMA concentration. Individual values are presented. There was 4-5 mice (except at 100 μM where it was 2 mice) at each of the 6 ADMA concentrations.** The K_m_ and V_max_ were calculated by Michaelis-Menten kinetic analysis of the total flux (individual values were used) which in this case is the same as saturable flux (GraphPad Prism version10). Unlabelled ADMA did statistically decrease the distribution of [^14^C]-sucrose into the pellet (S8 Fig). This suggests that the membrane integrity has not been compromised in these samples and could be presented here for comparison.

As the [^3^H]-ADMA concentration in the choroid plexuses, pituitary gland and pineal gland was higher than the concentration in the plasma at 10 minutes (i.e. R_Tissue_ > 100%; S12 Fig), single time uptake analysis could not be used to calculate the K_in_ needed to determine the total flux as described in the methods. Therefore, the K_m_, V_max_ and K_d_ values were not determined for these specific samples.

## DISCUSSION

The brain requires a constant supply of CAAs to function normally. CAAs are involved in normal brain cell growth, repair after damage and in regulating blood flow. The CAA, arginine, is the exclusive substrate for NO synthesis and the structurally related molecule, ADMA, is an inhibitor of NO production. Abnormal levels of CAA and their associated molecules (such as NO) are associated with several diseases including Alzheimer’s, a disease which is linked to hypoperfusion of brain tissues. Importantly, arginine has been used as a pre-training NO booster to increase blood flow in muscles during training, improve athletic performance and speed up recovery [15]. Arginine may protect against Alzheimer’s by increasing hippocampal NO levels [16]. Arginine supplementation has been shown to ameliorate the choroid plexus pathological changes in an Alzheimer’s disease aged rat model [17]. Interestingly, transporters for CAA can be used as a gateway for some medicines to enter the brain to treat brain disorders [18][19]. However, some cancer-causing viruses also use CAA transporters to get into the brain [20,21]. Consequently, furthering our understanding of the transport pathways of these two CAAs into the CNS is important and is relevant to several research areas due to the widespread influence of NO in physiological pathways in health and disease. The major contributors to the transport of both L-arginine and ADMA across the blood-CNS barriers were investigated as part of this present study, which builds on our previous work in this area [4][5].

Five CAAs transport systems have been identified according to their sensitivity to amino acids, inhibitors, and dependence upon Na^+^-gradients. These include the y^+^, y^+^L, b^o,+^, B^o,+^ and b^+^ transport systems [22][23][24]. Individual CAA may use more than one transport system. The term ‘transport system’ calls attention to the concept that a complex of different proteins rather than a single carrier protein mediates a distinct transport activity (S1 Table). Transporter systems for CAAs, which are expressed at the BBB include: y^+^, y^+^L, B^0,+^ and b^0,+^ systems [4,18,19,24–27].

We have previously demonstrated the presence of a saturable transport mechanism(s) at both the blood-brain and blood-CSF barriers for [^3^H]-arginine and [^3^H]-ADMA by means of the *in situ* brain/choroid plexus perfusion method in anaesthetized mice [5]. In this present study we investigated the identity of the transporters involved in this saturable transport of [^3^H]-arginine and [^3^H]-ADMA using the same method. To do this we included established inhibitors of specific amino acid transporters in the artificial plasma and compared results to those obtained in the absence of inhibitors (S1 Table). We first confirmed integrity of the blood-brain and blood-CSF barriers after exposure to the transporter inhibitors by checking that the values achieved for [^14^C]-sucrose were not significantly different in the absence and presence of the inhibitors (S1-S6 Fig). We then progressed to assessing the effect of inhibitors on the CNS delivery of [^3^H]-arginine and [^3^H]-ADMA. It was assumed that a significant change in the accumulation of [^3^H]-arginine or [^3^H]-ADMA into any CNS region was a result of a transporter interaction. A decrease in accumulation indicating the presence of an influx transporter and an increase in accumulation indicating the presence of an efflux transporter. Importantly, we noted that an absence of an inhibitor effect did not necessarily indicate that there was no significant transporter interaction. This is because transporter involvement in CAAs delivery to the CNS is difficult to interpret conclusively as: (i) CAA can be transported by both influx and efflux transporters plus some of the CAA transporters are bidirectional (e.g. system y^+^). (ii) CAA transporters are expressed on either the luminal and/or abluminal membranes of the brain capillary endothelial /choroid plexus epithelial cells. And (iii) [^3^H]-ADMA has been shown to be removed from the CNS [5].

In this present study we found that [^3^H]-arginine transport into all CNS regions (except CSF) was significantly inhibited (up to 99.7%) by *L*-homoarginine. This indicates that [^3^H]-arginine is entering into the CNS by system y^+^. This is because *L*-homoarginine is a substrate for system y^+^ [28] and the associated transporter proteins i.e. the cationic amino acid transporter-1 (CAT1; SLC7A1), CAT2A (SLC7A2) and CAT2B (SLC7A3) [29]. In addition, CAT1 is expressed at the BBB and the choroid plexus of the blood-CSF barrier [25][30][31][32]. Functionally, CAT1 is a bidirectional uniporter which facilitates the movement of CAAs down a concentration gradient and is usually involved in loading cells with amino acids [33]. System y^+^ activity has been demonstrated on both the luminal and abluminal plasma membranes of the bovine cerebral capillary endothelium, the site of the BBB [27]. Interestingly, our data suggests that system-y^+^ was present (at least) on the blood-side of the cerebral capillary endothelial cells and the choroid plexus due to reduced transport of [^3^H]-arginine into the cerebral capillary endothelial cell enriched pellet and the choroid plexus samples in the presence of *L*-homoarginine.

BCH and α-methyl-*D,L*-tryptophan are system B^0,+^ inhibitors, which can inhibit CAA (arginine) and neutral amino acid (leucine) transport by the system B^0,+^ transporter protein, ATB^0,+^ (SLC6A14) when expressed in xenopus oocytes or breast cancer cells [34][35] [36]. However, our *in situ* brain perfusion studies revealed that [^3^H]-arginine transport into the CNS was not affected by either of the system B^0,+^ inhibitors, BCH or α-methyl-*D,L*-tryptophan. Although it is noted that in one of the 15 CNS regions (i.e. hippocampus) there was a decrease in accumulation (i.e. 47.5%) observed with one (but not both) of the inhibitors (i.e. α-methyl-*D,L*-tryptophan). It is possible that this is just a sampling error but may reflect specific inhibitor sensitivity in this region.

Overall, our data provides evidence that [^3^H]-arginine predominately uses system-y^+^, and not system-B^0,+^, to cross both the BBB and the blood-CSF barrier (choroid plexus). This agrees with earlier studies by us and others which showed that: i) transporters for [^3^H]-arginine could be detected on the luminal membrane of the cerebral capillary endothelium and the blood-side of the choroid plexus [5][27]. ii) arginine can inhibit the cellular uptake of *L*-homoarginine by system-y^+^ transporter proteins such as CAT1 [29]. iii) [^3^H]-arginine transport at the human BBB *in vitro* was not affected by the system B^0,+^ substrate, leucine [4]. iv) arginine transport by the transporter protein for system B^0,+^, ATB^0,+^, may not be detectable due to a preference for system y^+^ and/or the presence of other system B^0,+^ substrates [37] and/or the low expression of ATB^0,+^ in normal tissues [36]. v) the transporter for arginine at the blood-side of the isolated perfused choroid plexus was found not to be sensitive to the system-B^0,+^ inhibitor, BCH [38] and vi) sodium-independent influx of arginine at physiological concentrations occurs predominately by a single and saturable transport system [28].

The next part of our study was to investigate the identity of the transporters involved in the saturable [^3^H]-ADMA transport across the BBB and the blood-CSF barriers [5]. The significant inhibition of [^3^H]-ADMA uptake into all the brain regions sampled (up to 99.2%) and the choroid plexus (88.2%) in the presence of 20 mM *L*-homoarginine would suggest that [^3^H]-ADMA transport at the BBB and blood-CSF barrier can be mediated by system-y^+^ (Figures 4 and 6; S2 and S4 Tables). The transport of [^3^H]-ADMA into the circumventricular organs (the pineal and pituitary glands) was also sensitive to *L*-homoarginine indicating system y^+^-involvement (Figure 6). Other studies have also provided evidence that ADMA transport may be sensitive to *L*-homoarginine, as ADMA inhibited *L*-homoarginine uptake into human embryonic kidney (HEK293) cell lines stably overexpressing CAT1 [29]. In addition, we have previously shown that [^3^H]-arginine transport across the BBB and blood-CSF barrier is inhibited by ADMA and *vice versa* [5] and in this present study we have shown that [^3^H]-arginine appears to use system y^+^. Together this information supports a conclusion that [^3^H]-ADMA transport across the BBB and the blood-CSF barrier involves system-y^+^ [5].

However, as the uptake of [^3^H]-ADMA into certain brain regions and the choroid plexus was also significantly inhibited (albeit to a lesser degree) by the presence of the other transporter inhibitors we have evidence that other transporter systems are involved, in addition to system y^+^ (S2 and S4 Tables). Interestingly, our results indicated that [^3^H]-ADMA uptake into the brain and choroid plexus was inhibited up to 50.0% by the two system B^0,+^ inhibitors (BCH and α-methyl-*D,L*-tryptophan). Therefore, ADMA is likely to be also interacting with system B^0,+^ for transport from plasma to the brain via the BBB and plasma to CSF via the choroid plexus. This is also supported by the fact that [^3^H]-ADMA delivery to the CNS was inhibited by leucine. Leucine is a neutral amino acid which is not transported by system-y^+^ [39], but is a substrate for system B^0,+^. In addition, NOS inhibitors, which, like ADMA, are structurally related to arginine (for example N^G^-nitro-L-arginine), have also been shown to be transported by ATB^0,+^ [40]. We, and others, have previously identified the presence of ATB^0,+^ in human BBB cells (hCMEC/D3) using Western blotting and immunofluorescence [4][19][18]. It is thought to be a sodium-dependent symporter involved in the uptake of amino acids into BBB cells (hCMEC/D3) [18,33] [4][19]. Together these facts provide further support that ADMA could also be transported by system B^0,+^ (ATB^0,+^) across the brain capillary endothelial cells and choroid plexus epithelium. Although, it is noted that ATB^0,+^ mRNA has not been detected in human choroid plexus samples [41].

Interestingly, as well as being transported by ATB^0,+^, leucine and BCH are substrates for system-*L* [42]. System *L*-amino acid transporters such as the large neutral amino acid transporter 1 protein (LAT1) are encoded by the SLC7A5 gene and forms a heterodimer with the glycoprotein CD98, which is encoded by the SLC3A2 gene. System-*L* transporters have been shown to be expressed on both luminal and abluminal membranes of the cerebral endothelial cell and the basolateral surface (blood-side) and apical side (CSF-side) of the choroid plexus [43][44][45][24][46]. LAT1 is a sodium-independent high-affinity transporter for many of the neutral amino acids and prefers those with a bulky side chain, such as leucine and phenylalanine. Both leucine and phenylalanine transport at the choroid plexus has been shown to be sensitive to BCH [43]. Interestingly, the L-system carrier at the choroid plexus is thought to be involved in the transport of leucine and phenylalanine from blood to CSF and CSF to blood [45]. Interestingly, our data also show that [^3^H]-ADMA transport into some regions of the brain and the choroid plexus are also sensitive to phenylalanine with uptake being inhibited by up to 66.4%. Thus as ADMA is positively charged and will not be being transported by the neutral amino acid transporter, system L, our data suggests that phenylalanine is interacting with system B^0,+^. Our data would confirm that system B^0,+^ has a broad substrate selectivity (denoted by ‘B’) accepting neutral (denoted by ‘0’) and cationic (denoted by ‘+’) amino acids as substrates.

Other CAA transport systems may also be involved in [^3^H]-ADMA delivery to the CNS. For example, system y^+^L and system b^0,+^. System y^+^L is a heterodimer formed by the interaction of the heavy chain of the cell surface antigen 4F2 (4F2hc, encoded by the SLC3A2gene) with the light chains 4F2-lc2 (or y^+^LAT-1, encoded by SLC7A7) or 4F2-lc3 (or y^+^LAT-2, encoded by SLC7A6). System y^+^L transports CAAs with no need for extracellular Na^+^; however, it can transport both small and large neutral amino acids with high affinity in the presence of this cation [47]. In fact, system y^+^L transporters such as y^+^LAT1 and y^+^LAT2 are thought to exchange a neutral amino acid coupled to a sodium ion for a CAA. System y^+^L transporters are expressed at the human BBB e,g. y^+^LAT1 (SLC7A7) and y^+^LAT2 (SLC7A6) [24]. SLC7A7, SLC7A6 and SLC3A2 mRNA has also been detected in human choroid plexus[31,48][49][50]. In the presence of a CAA loader, CAT1, y^+^LAT1 mediates efflux of CAAs in exchange for neutral amino acids and so these amino acid transporters exhibit functional co-operation. The fact that leucine and phenylalanine inhibited [^3^H]-ADMA accumulation could also be explained by stimulation of ADMA efflux via system y^+^L by these neutral amino acids. As system y^+^L removes CAA from cells, this transporter may be involved in the removal of ADMA from the CNS. Removal of ADMA from the CNS has previously been observed by our group [5].

System b^0,+^ is a facilitated transporter of CAAs, which is inhibited by the organic cation, harmaline [27]. It transports cationic and zwitterionic amino acids. The transporter protein associated with system b^0,+^ is b^0,+^AT and is encoded by the SLC7A9 gene. b^0,+^AT mRNA is expressed in the human brain capillary endothelial cells, but at low levels [24] and b^0,+^AT mRNA was not found in human choroid plexus samples [51]. However, harmaline significantly inhibited [^3^H]-ADMA uptake into the brain and choroid plexus suggesting the involvement of system b^0,+^ in ADMA transport across the mouse BBB and blood-CSF interfaces. The conflicting results may reflect species differences in transporter expression and/or lack of inhibitor specificity for the system b^0,+^ transporter.

Interestingly, [^3^H]-ADMA accumulation into the brain regions was more sensitive to the transporter inhibitor, L-homoarginine, than unlabelled ADMA (Figure 4; S10 Fig; S2 Table). For example, 20 mM L-homoarginine decreased the uptake of [^3^H]-ADMA by 95.7%, whereas 500 μM unlabelled ADMA only decreased the uptake of [^3^H]-ADMA by 79.9%, into the frontal cortex. This could be related to concentration differences, but may be because [^3^H]-ADMA delivery to the CNS involves multiple cationic amino acid transporters such as system y^+^, B^0,+^, y^+^L and b^0,+^. As these transporters are involved in the influx and efflux of ADMA, all could be affected to some degree by the presence of unlabelled ADMA and as a result the overall uptake into the brain of ADMA in the presence of unlabelled ADMA could be smaller than that achieved with the specific inhibitor, L-homoarginine.

In this present study we also determined the K_m_ (half-saturation constant), V_max_ (maximal influx rate of the saturable component) and K_d_ (diffusion constant) for [^3^H]-ADMA transport into the different CNS regions using an ADMA concentration range of 62.5 nM – 500 μM at a single time point of 10 minutes (S5 Table). For [^3^H]-ADMA delivery into the frontal cortex the K_m_, V_max_ and K_d_ were 29.07±7.19 μM, 0.307±0.017 nmol.min^-1^.g^-1^ and 4.40 μl.min^-1^.g^-1^ respectively. This compares with [^3^H]-ADMA delivery into the brain homogenate having a K_m_ of 13.59±5.26 μM, V_max_ of 0.155±0.013 nmol.min^-1^.g^-1^ and K_d_ of 3.34 μl.min^-1^.g^-1^, into the brain supernatant having a K_m_ of 68.52±44.41 μM, V_max_ of 0.208±0.036 nmol.min^-1^.g^-1^ and K_d_ of 2.35 μl.min^-1^.g^-1^ and into the pellet of a K_m_ of 33.97±16.83 μM, V_max_ of 0.071±0.008 nmol.min^-1^.g^-1^ and K_d_ of 0.38 μl.min^-1^.g^-1.^ The K_m_ and V_max_ values for [^3^H]-ADMA accumulation into the CSF being 30.59±25.41 μM, and 2.07±0.38 nmol.min^-1^.g^-1^, respectively. We could not detect a non-saturable component to the total flux for [^3^H]-ADMA delivery into the CSF, so were not able to calculate a diffusion constant (K_d_) for this region. As we have evidence that ADMA uses multiple transporters to cross the BBB and blood-CSF barrier and can cross from blood-to-CNS and CNS to blood, it is not possible to assign the kinetic constants to a specific transporter.

The plasma concentration of ADMA is 0.30-1.57 μM in humans [13] and 1.07-1.58 μM in mice [52]. Plasma ADMA concentrations are raised in diabetes mellitus [53], hypertension [54] and stroke [55], reaching approximately 3 μM in certain disease states [13]. Interestingly, it is also significantly elevated after prolonged, strenuous exercise (0.32±0.05 versus 0.37±0.05 μM) possibly as a result of exercise-induced muscle damage and inflammation [56]. This suggests that the saturable transport system(s) for [^3^H]-ADMA delivery into the CNS identified in this study would not be fully saturated by ADMA in the plasma in health or in disease. This may be as expected as a single CAA transport system will transport several CAAs.

Interestingly, our previous study revealed that ADMA and arginine shared a transporter to some extent [5] and the transporter inhibitor studies described in this present study confirmed that arginine and ADMA are transported by system y^+^. It is interesting to note that system y^+^ usually binds substrates with a relatively high affinity (K_m_<200μM) [28] and CAT-1 transports ADMA with a K_m_ of 183±21 μM [57]. In addition, system y^+^ is a bidirectional uniporter mediating the influx and efflux of positively charged amino acids with the K_m_ for influx being lower than the K_m_ for efflux. This asymmetry may arise in part from the transmembrane potential [28]. The transporter kinetics (Table S5) and plasma concentrations described in this present study and our previous observations [5] would allow arginine supplementation to increase NO production by reducing the intracellular concentration of ADMA either by competitively inhibiting ADMA influx and/or by stimulating ADMA efflux. Thus transporter interaction would explain the arginine paradox, whereby nutritional supplementation with arginine increases NO production despite the intracellular NOS being saturated with arginine. Although other mechanisms such as changes to ADMA catabolism [58] and NOS affinity [13][59] cannot be ruled out.

The advantages and limits of the *in situ* brain perfusion method have recently been described by us [5]. In this present study a further advantage is demonstrated and that is the ability to manipulate the composition of the artificial plasma to measure the kinetic constants (K_m_, V_max_ and K_d_) of test molecule (in this case ADMA) transport. A limit of this is that these constants do not necessarily reflect those of a single transporter, but represent blood-CNS (and possibly CNS-blood) transport of the molecule, which could be the result of multiple transporter interactions. A confounding factor of the present study was the actual significance of the statistical difference observed with [^14^C]-sucrose between the control and some of the test groups. In particular when the [^14^C]-sucrose values overlapped with other groups where no statistical difference was detected. We evaluated the relevance of these differences carefully and on a case by case level. Importantly, if we remove these groups of data our conclusions still remain the same.

## Conclusion

The results of our present study indicate that [^3^H]-arginine transport at the BBB and the blood-CSF barrier (choroid plexus) involves the cationic amino acid transport system, system y^+^, but not system B^0,+^. Our study also provides new information that [^3^H]-ADMA delivery to the CNS is more complex and involves multiple CAA transporters including system y^+^, system B^0,+^, system y^+^L and system b^0,+^. Overall, this suggests that the intracellular concentration of ADMA is under tight control as multiple pathways are involved in its regulation and this is likely related to its ability to inhibit NO production. A major contributor to the transport of both *L*-arginine and ADMA at both the BBB and BCSFB is the bidirectional transport system, system y^+^. As system y^+^ is also a transport system that removes cationic amino acids from cells, it is plausible that this particular transport system, together with the y^+^L -system, is involved in the CNS to blood efflux of ADMA that we have previously observed [5]. In this present study we also determined the kinetic constants of [^3^H]-ADMA delivery into the CNS from the blood. Results suggested that the transporters for ADMA at the BBB and blood-CSF barrier would not be completely saturated by ADMA at the concentrations normally found in the plasma. This would allow these transporters to be utilized by other endogenous cationic amino acids that are needed by the brain to function normally. It remains plausible that the positive effects of arginine supplementation are due to changes in intracellular ADMA concentrations in part as a direct result of transporter interaction (likely system y^+^) and this can help explain the arginine paradox. Arginine supplementation in diseases such as AD and stroke is an interesting strategy to improve blood flow and reduce endothelial dysfunction.

## DATA SHARING

All data underlying the results are available as part of the article and no additional source data are required.

## FUNDING

This work was supported by a Biotechnology and Biological Sciences Research Council (BBSRC) centre for integrative biomedicine PhD studentship for Mr Fidanboylu to Dr Sarah Ann Thomas [BB/E527098/1]. https://www.ukri.org/councils/bbsrc/. This research was funded in whole, or in part, by the Wellcome Trust [080268]. https://wellcome.org/. The recipient of this grant was Dr Sarah Ann Thomas. For the purpose of Open Access, the author has applied a CC BY public copyright licence to any Author Accepted Manuscript version arising from this submission. The funders had no role in study design, data collection and analysis, decision to publish, or preparation of the manuscript.

## CONFLICT OF INTEREST

The authors declare that the research was conducted in the absence of any commercial or financial relationships that could be construed as a potential conflict of interest.

## Abbreviations

ADMA: asymmetric dimethylarginine or N^G^,N^G^-dimethyl-*L*-arginine
BCH: 2-amino-endo-bicyclo[2.2.1]heptane-2-carboxylic acid
BBB: blood-brain barrier
CAAs: cationic amino acids
CAT: cationic amino acid transporter
CNS: central nervous system
CSF: cerebrospinal fluid
CVO: circumventricular organ
K_d_: constant of non-saturable diffusion
eNOS: endothelial nitric oxide synthase
K_m_: half-saturation constant
*V_max_*: maximal influx rate of the saturable component
C_cap_: , mean capillary concentration of the amino acid
NO: nitric oxide
NOS: nitric oxide synthase
K_in_: unidirectional transfer constant
J_in_: unidirectional flux
SD: standard deviation
SEM: standard error of the mean

## ACKNOWLEDGEMENTS

This paper includes data from the PhD thesis of Mehmet Fidanboylu [60]. Abstracts of this work have been published [61].

## Author contributions

**MF:** Conceptualization, Data Curation; Validation, Formal Analysis, Investigation, Writing – Review & Editing, Project Administration, Visualization

**SAT**: Conceptualization, Data Curation; Validation, Funding Acquisition; Investigation, Formal Analysis, Supervision, Visualization, Writing – Original Draft Preparation, Writing – Review & Editing, Resources, Project Administration

## Supplementary information files

**S1 Fig: The effect of *L*-homoarginine, BCH and α-methyl-*D*,*L*-tryptophan on the regional brain uptake of [^14^C]-sucrose (10 minute perfusion; co-perfused with [^3^H]arginine).** Uptake is expressed as the percentage ratio of tissue to plasma (mL.100 g^-1^). Each bar represents the mean ± SEM of 3-6 animals Each marker represents one animal. One-way ANOVA with Dunnett’s post-hoc test was used to compare means to control ([^14^C]-sucrose alone), with statistical significance taken as \**p* <0.05, **p<0.01, ***p<0.001 (GraphPad Prism 10.2 for Windows).

**S2 Fig: The effect of *L*-homoarginine, BCH and α-methyl-*D*,*L*-tryptophan uptake of [^14^C]-sucrose in capillary depletion samples (10 minute perfusion; co-perfused with [^3^H]arginine).** Uptake is expressed as the percentage ratio of tissue to plasma (mL.100 g^-1^). Each bar represents the mean ± SEM of 4-6 animals. Each marker represents one animal. One-way ANOVA with Dunnett’s post-hoc test was used to compare means to control ([^14^C]-sucrose alone), with statistical significance taken as \**p* <0.05, **p<0.01, ***p<0.001 (GraphPad Prism 10.2 for Windows).

**S3 Fig: The effect of *L*-homoarginine, BCH and α-methyl-*D*,*L*-tryptophan on the uptake of [^14^C]-sucrose in CSF, choroid plexus and pituitary gland samples (10 minute perfusion; co-perfused with [^3^H]arginine).** Uptake is expressed as the percentage ratio of tissue to plasma (mL.100 g^-1^). Each bar represents the mean ± SEM of 4-6 animals, except the CSF samples for the *L*-homoarginine and the α-methyl-*D*,*L*-tryptophan inhibitor groups where n=3 and n=2, respectively. Each marker represents one animal. One-way ANOVA with Dunnett’s post-hoc test was used to compare means to control ([^14^C]-sucrose alone), with statistical significance taken as \**p* <0.05, **p<0.01, ***p<0.001 (GraphPad Prism 10.2 for Windows).

**S4 Fig: The effect of 20 mM *L*-homoarginine, 4 mM BCH, 500 μM a-methyl-*D*,*L*-tryptophan, 200 μM *L-*phenylalanine, 5 mM *L*-leucine and 2 mM harmaline on the regional brain uptake of [^14^C]-sucrose (10 minute perfusion; co-perfused with [^3^H]ADMA).** Uptake is expressed as the percentage ratio of tissue to plasma (mL.100 g^-1^). Perfusion time is 10 minutes. Each bar represents the mean ± SEM of 4-5 animals. Each marker represents one animal. Asterisks represent one-way ANOVA with Dunnett’s post-hoc tests comparing mean±SEM to control within each sample/region, \**p* < 0.05, \*\**p* < 0.01, \*\*\**p* < 0.001 (GraphPad Prism 10.2 for Windows).

**S5 Fig: The effect of 20 mM *L*-homoarginine, 4 mM BCH, 500 μM a-methyl-*D*,*L*-tryptophan, 200 μM *L-*phenylalanine, 5 mM *L*-leucine and 2 mM harmaline on the distribution of [^14^C]-sucrose in capillary depletion samples (10 minute perfusion; co-perfused with [^3^H]-ADMA).** Uptake is expressed as the percentage ratio of tissue to plasma (mL.100 g^-1^) and is corrected for [^14^C]-sucrose. Each marker represents one animal. Perfusion time is 10 minutes. Each bar represents the mean ± SEM of 4-5 animals. Asterisks represent one-way ANOVA with Dunnett’s post-hoc tests comparing mean±SEM to control within each region/sample, \**p* < 0.05, \*\**p* < 0.01 (GraphPad Prism 10.2 for Windows).

**S6 Fig: The effect of 20 mM *L*-homoarginine, 4 mM BCH, 500 μM a-methyl-*D*,*L*-tryptophan, 200 μM *L-*phenylalanine, 5 mM *L*-leucine and 2 mM harmaline on the distribution of [^14^C]-sucrose in the CSF and CVOs (10 minute perfusion; co-perfused with [^3^H]-ADMA).** Uptake is expressed as the percentage ratio of tissue to plasma (mL.100 g^-1^) and is corrected for [^14^C]-sucrose. Perfusion time is 10 minutes. Each marker represents one animal. Each bar represents the mean ± SEM of 4-5 animals. Asterisks represent one-way ANOVA with Dunnett’s post-hoc tests comparing mean±SEM to control within each region/sample, \**p* < 0.05, \*\**p* < 0.01 (GraphPad Prism 10.2 for Windows).

**S7 Fig: The effect of 0.5, 3, 10, 100 and 500 μM un-labelled ADMA on the uptake of [**^14^**C]-sucrose in the brain (10 minute perfusion; co-perfused with [^3^H]-ADMA).** Uptake is expressed as the percentage ratio of tissue to plasma (mL.100 g^-1^). Perfusion time is 10 minutes. Each bar represents the mean ± SEM of 4-5 animals. Each marker represents one animal. Asterisks represent one-way ANOVA with Dunnett’s post-hoc tests comparing mean±SEM to control, \**p* < 0.05, \*\**p* < 0.01, \*\*\**p* < 0.001 (GraphPad Prism version 10.2.2 for windows).

**S8 Fig: The effect of 0.5, 3, 10, 100 and 500 μM un-labelled ADMA on the uptake of [**^14^**C]-sucrose in the capillary depletion samples (10 minute perfusion; co-perfused with [^3^H]-ADMA).** Uptake is expressed as the percentage ratio of tissue to plasma (mL.100 g^-1^). Perfusion time is 10 minutes. Each bar represents the mean ± SEM of 4-5 animals. Each marker represents one animal. Asterisks represent one-way ANOVA with Dunnett’s post-hoc tests comparing mean±SEM to control, \**p* < 0.05, \*\**p* < 0.01, \*\*\**p* < 0.001 (GraphPad Prism version 10.2.2 for windows). The [^14^C]-sucrose distribution into the pellet samples was statistically reduced by the presence of unlabelled ADMA at most concentrations, but was in the range achieved in other test groups where no statistical difference was obtained (S2 Fig). This suggests the membrane was intact in these samples.

**S9 Fig: The effect of 0.5, 3, 10, 100 and 500 μM un-labelled ADMA on the uptake of [**^14^**C]-sucrose in the CSF and CVOs (10 minute perfusion; co-perfused with [^3^H]-ADMA).** Uptake is expressed as the percentage ratio of tissue to plasma (mL.100 g^-1^). Perfusion time is 10 minutes. Each bar represents the mean ± SEM of 3-5 animals. Each marker represents one animal. Asterisks represent one-way ANOVA with Dunnett’s post-hoc tests comparing mean±SEM to control, \**p* < 0.05, \*\**p* < 0.01, \*\*\**p* < 0.001 (GraphPad Prism version 10.2.2 for windows).

**S10 Fig: The effect of 0.5, 3, 10, 100 and 500 μM un-labelled ADMA on the uptake of [**^3^**H]-ADMA in the brain.** Uptake is expressed as the percentage ratio of tissue to plasma (mL.100 g^-1^) and is corrected for [^14^C]-sucrose (vascular space). Perfusion time is 10 minutes. Each bar represents the mean ± SEM of 4-5 animals. Each marker represents one animal. Asterisks represent one-way ANOVA with Dunnett’s post-hoc tests comparing mean±SEM to control, \**p* < 0.05, \*\**p* < 0.01, \*\*\**p* < 0.001 (GraphPad Prism 6.0 for Mac).

**S11 Fig: The effect of 0.5, 3, 10, 100 and 500 μM un-labelled ADMA on the distribution of [^3^H]-ADMA in capillary depletion samples.** Uptake is expressed as the percentage ratio of tissue to plasma (mL.100 g^-1^) and is corrected for [^14^C]-sucrose. Each bar represents the mean ± SEM of 4-5 animals. Each marker represents one animal. Asterisks represent one-way ANOVA with Dunnett’s post-hoc tests comparing mean±SEM to control, \**p* < 0.05, \*\**p* < 0.01, \*\*\**p* < 0.001 (GraphPad Prism version 10.2.2 for windows).

**S12 Fig: The effect of 0.5, 3, 10, 100 and 500 μM unlabelled ADMA on the distribution of [**^3^**H]-ADMA in the CSF and CVOs.** Uptake is expressed as the percentage ratio of tissue or CSF to plasma (mL.100 g^-1^) and is corrected for [^14^C]-sucrose. Each bar represents the mean ± SEM. n= 4-5 mice (for the CSF samples except at 100 μM where it was 2 mice), 4-5 mice (for the choroid plexus samples), 5 mice (for the pineal gland samples except at 100 μM where it was 3 mice) and 4-5 mice (for the pituitary gland samples) at each of the 6 ADMA concentrations. (GraphPad Prism version 10.2.2 for windows). Asterisks represent one-way ANOVA with Dunnett’s post-hoc tests comparing mean±SEM to control, \**p* < 0.05, \*\**p* < 0.01, \*\*\**p* < 0.001.

**S13 Fig: The contributions of the saturable and non-saturable components to total brain flux of [^3^H]ADMA are plotted against the unlabelled ADMA concentration.** The measured values are the mean ±SEM for 4-5 mice at each of the 6 ADMA concentrations and has been [^14^C]-sucrose corrected. The lower lines show the contributions of the saturable and non-saturable components to total influx. The K_d_ value was calculated from linear regression analysis of the total flux at the highest concentrations. The K_m_ and V_max_ were calculated by Michaelis-Menten kinetic analysis of the saturable flux (mean values were used). Analyses were performed using GraphPad Prism version10. Unlabelled ADMA concentrations of 500 μM did statistically increase the distribution of [^14^C]-sucrose into the occipital cortex, caudate nucleus, hippocampus, thalamus, pons and cerebellum and the values measured were at the upper range of [^14^C]-sucrose values achieved at the lower concentrations of unlabelled ADMA (i.e. ≤100 μM unlabelled ADMA; S7 Fig). As this does suggest loss of BBB integrity in these regions, we only interpreted the kinetic characteristics of [^3^H]-ADMA in the other regions where the BBB remained statistically intact (i.e. frontal cortex, hypothalamus). However, all regions are presented here for comparison.

**S1 Table: The amino acid transport systems studied, the transporter inhibitors, the inhibitor concentration utilised, the transporter protein inhibited and their respective gene codes according to the human genome organisation (HUGO).** A complex of different proteins may mediate a distinct system activity.

**S2 Table: Results obtained from specific transport-inhibition studies for [^3^H]-ADMA uptake into the brain.** Uptake is expressed as the percentage ratio of tissue to plasma (mL.100 g^-1^) and is corrected for [^14^C]-sucrose (vascular space). Perfusion time is 10 minutes. One-way ANOVA with Dunnett’s post-hoc test was used to compare means to control ([^3^H]-ADMA alone), with statistical significance taken as *p* < 0.05. The percentage inhibition is reported where appropriate. Each value has been obtained from an n of 5 experiments except where stated.

**S3 Table: Results obtained from specific transport-inhibition studies for [^3^H]-ADMA in capillary depletion samples.** Uptake is expressed as the percentage ratio of tissue to plasma (mL.100 g^-1^) and is corrected for [^14^C]-sucrose. One-way ANOVA with Dunnett’s post-hoc test was used to compare means to control ([^3^H]-ADMA alone), with statistical significance taken as *p* < 0.05. n = number of experiments. The percentage inhibition is reported where appropriate. Each value has been obtained from an n of 5 experiments except where stated. n.s. = not significant. #The [^14^C]-sucrose value in that compartment was significantly increased by the presence of the inhibitor, *L-*leucine.

**S4 Table: Results obtained from specific transport-inhibition studies for [^3^H]-ADMA in the CSF, choroid plexus and CVO’s.** One-way ANOVA with Dunnett’s post-hoc test was used to compare means to control ([^3^H]-ADMA alone), with statistical significance taken as *p* < 0.05. n = number of experiments. Each value has been obtained from an n of 5 experiments except where stated.

**S5 Table: The kinetic constants for [^3^H]-ADMA influx into the different brain regions, capillary depletion samples and the CSF.** The K_m_ and V_max_ values were calculated using the saturable flux, which was determined from the total flux by correcting for non-saturable flux (K_d_ × concentration). Except those for CSF which were calculated from the total flux as there was no non-saturable component. **^#^**The [^14^C]-sucrose (vascular space) in the majority of brain region (except the frontal cortex and hypothalamus) was significantly increased by the presence of the 500 μM unlabelled ADMA. Despite this the kinetic constants (K_m_) values calculated in the each of the brain regions were not statistically significantly different to each other (One-Way ANOVA followed by Tukey’s multiple comparison test). ^†^The [^14^C]-sucrose distribution into the pellet samples was statistically reduced by the presence of unlabelled ADMA at all concentrations (except 100 μM ADMA where there was no difference) but was in the range achieved in other test groups where no statistical difference was obtained (S2 Fig). This suggests the membrane was intact in these samples.

**S1 Data Set:** [^3^H]-arginine and [^14^C]-sucrose in the absence and presence of transporter inhibitors.

**S2 Data Set**: [^3^H]-ADMA and [^14^C]-sucrose in the absence and presence of transporter inhibitors.

**S3 Data Set:** [^3^H]-ADMA and [^14^C]sucrose in the absence and presence of unlabelled ADMA at various concentrations.

## Notes

### Competing Interest Statement

The authors have declared no competing interest.

